# Susagi: A Microbiome World Model

**DOI:** 10.64898/2026.05.07.723428

**Authors:** Matteo Peluso, Janko Tackmann, Christian von Mering

## Abstract

**Motivation:** Accurately modelling how microbial communities assemble and change across hosts and environments is essential for analysis and intervention. Typical pipelines capture limited generalisable structure and often depend on fixed ecological unit definitions.

**Results:** We present Susagi (Set Unsupervised Assessment of Genetic Imposters), a permutation-invariant denoising transformer that operates directly on sets of bacterial SSU rRNA gene embeddings to learn a member-level *stability* function.

The model was trained on ∼ 2 × 10^6^ bacterial community samples. We show that it reliably predicts community composition dynamics in a zero-shot (no training) setting, demonstrated here across three challenging microbiomes for which traditional ML methods do not exceed random expectation. The model’s stability scores capture biological structure: across datasets, higher scores are enriched for agricultural, cropland, and soil-associated habitats, consistent with microbial communities in these environments supporting positive diversity–stability relationships. The highest stability scores are only attained by communities with high Pielou evenness and large size, despite the fact that the model has never seen abundances suggesting it can recognise community dysbiosis from presence absence alone. Furthermore, they also track biological gradients such as subject age.

Susagi is competitive with another Large Microbiome Model (Microbial General Model, MGM) on diverse classification tasks, without task-specific fine-tuning and with an increased parameter efficiency.

Ultimately, our model will facilitate hypothesis generation for complex microbial processes, including deterministic assembly and microbial interactions, crucial for instance in the design of communities *in silico*.

**Availability and implementation:** Evaluation code and model weights can be found from https://github.com/the-puzzler/Microbiome-Modelling. Model weights can also be downloaded directly from https://huggingface.co/basilboy/microbiome-model. Interactive demo can be found here: https://huggingface.co/spaces/basilboy/microbiome-space.

## 1 Introduction

Microbial communities shape ecosystems and host biology by mediating nutrient cycling, pathogen exclusion, and responses to environmental change [1–3]. Microbes in these communities are not independent units, but rather exist in complex interacting structures [4]. Untangling these complex relationships is challenging and resource-intensive in the laboratory. Computational tools that attempt to implicitly learn the underlying community structures from experimental data could help to narrow the search space of experiments by providing a data-backed *in silico* test bed for hypothesis generation.

Large-scale SSU rRNA resources with harmonized metadata now make it feasible to learn shared structure across diverse microbiomes spanning environments, hosts and conditions. [5–8]. Historically, predictive pipelines have either treated communities as bag-of-OTUs inputs for sample-level classifiers or summarised dependence structure via network approaches (often utilising co-occurrence), which typically only indirectly captures the higher-order, context-dependent interactions thought to underlie assembly and stability processes. More recently, deep sequence models, graph neural networks, and neural dynamical systems have started to model microbiome structure and dynamics more holistically [9–13]. In practice, however, these deep models are typically instantiated as study-specific predictors: for example, graph neural networks trained per wastewater treatment plant, recurrent networks fit to individual bioreactor experiments, and sequence or profile “language models” optimized within particular cohorts. They generally rely on fixed ecological unit definitions tied to reference databases [9–12]. Even methods trained on global microbiome datasets usually learn embeddings for a fixed set of OTUs, rather than explicitly leveraging the idea that SSU rRNA sequences themselves encode broader evolutionary, functional and ecological information [14]. As summarized by Przymus et al. [13], this ecosystem of methods has substantially advanced within-study prediction and control, but still falls short of providing a universal, sequence-based descriptive language that can be applied consistently across diverse microbiome types. Furthermore, training on specific datasets has increased risk of learning artifacts that reduce practical reusability.

Permutation-invariant neural architectures (e.g. Transformers used without positional encodings, Deep Sets, Set Transformer, graph neural networks) naturally operate on unordered collections and can capture interactions without imposing arbitrary sequence order [15–18]. In parallel, DNA language models provide informative, transferable and threshold independent representations for short nucleotide sequences, including SSU rRNA segments [19, 20]. Prior microbiome models typically order OTUs by abundance, and use fixed ecological unit definitions based on rigid taxonomies. This limits open vocabulary generalisation, relies on noisy relative read counts and neglects the potential of pre-learned DNA sequence embeddings as rich, generalisable semantic units [12, 14].

To overcome these restrictions we introduce Susagi (Set Unsupervised Assessment of Genetic Imposters), a permutation-invariant denoising transformer that estimates a member-level *community-stability* function: given a set of microbial SSU rRNA embeddings (and optional text metadata), the model assigns each candidate a logit indicating whether it plausibly belongs to the community under the given context. This is shown in **Training Process** of Fig. 1. Intuitively, this community-stability function reflects which OTUs are compatible and persistent given the rest of the community and its metadata. Our framing is loosely analogous to protein language models, which treat naturally occurring amino-acid sequences as the outcome of evolutionary selection and use logits over substitutions to infer which mutations are compatible with a stable fold or function [21]. Here, we treat observed environmental microbiomes as approximate “stable states” of microbial communities and train the model to distinguish resident OTUs from nearby imposters in SSU-rRNA-embedding space, with the goal of encoding fundamental member-level, and by extension sample-level, stability in the resulting logits [6].

**Figure 1:**
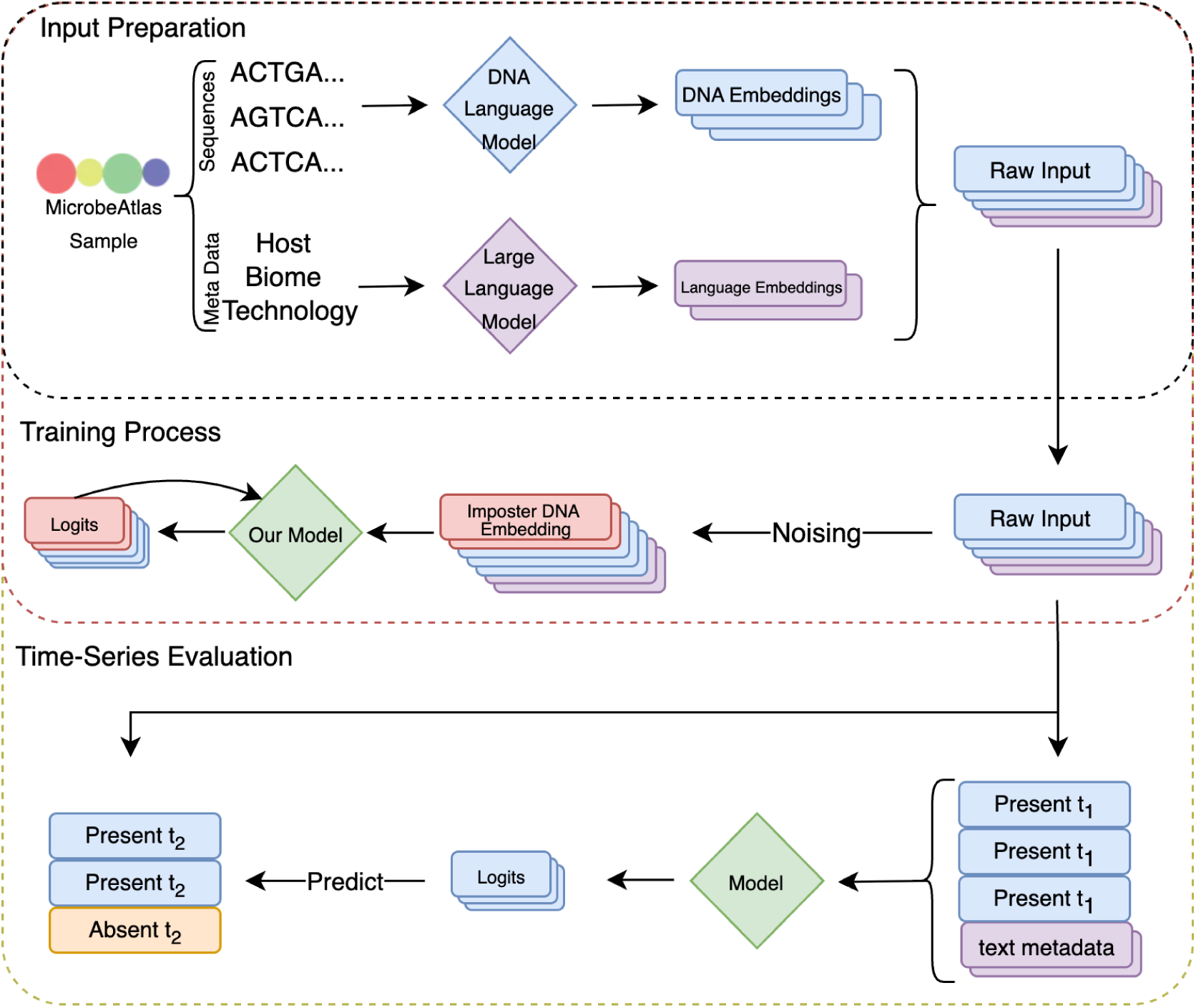
Imposter training for microbiome predictions. **Input Preparation:** inputs are a set of SSU rRNA gene embeddings and optional text embeddings for metadata. **Training Process:** we create negative (imposter) inputs by adding OTUs from a pool in DNA embedding space defined by minimum similarity. A permutation-invariant transformer outputs per-token logits (stability scores) indicating true resident vs. added noise. **Time series evaluations:** time series evaluations are performed by passing unseen raw samples through the model and comparing the output to presence/absence patterns at subsequent time points.

In summary, we (i) formalize community stability as permutation-invariant set prediction, avoiding noisy relative-abundance orderings, and introduce a denoising-set objective with negatives created via minimum-similarity pool sampling; (ii) show that operating directly in SSU-rRNA-embedding space yields parameter-efficient models that can score previously unseen sequences; (iii) demonstrate zero-shot temporal generalization across diverse and challenging environments; (iv) show that the frozen sample encoder provides transferable set-level representations for downstream tasks; (v) under a controlled, lightweight protocol (frozen encoder plus linear heads), achieve performance comparable to a more parameter heavy, fine-tuned state-of-the-art model (the Microbial General Model, MGM); and (vi) theorise that the shown prediction capacity originates from the formation of a latent community-space with local attractors under the training objective.

## 2 Materials and Methods

### 2.1 Datasets

#### Training corpus

We downloaded ∼ 2×10^6^ microbiome sequencing samples (presence/absence for this work) with harmonized metadata from the MicrobeAtlas database [5, 6]. It covers human-, animal-, plant-associated as well as various environmental sources. We excluded all samples belonging to the induced gingivitis [22], snowmelt [23], and DIABIMMUNE [24, 25] studies, so that no subject or time point from these datasets was seen during pretraining. The infant cohort [26] used for downstream classification is not part of MicrobeAtlas. The IBS cohorts used for classification are included in MicrobeAtlas; in this setting we allow overlap because we explicitly train supervised linear heads on top of the frozen encoder and the metadata do not contain the disease labels.

#### Induced gingivitis (zero-shot temporal weekly) [22]

Longitudinal oral time series during induced, then alleviated gingivitis. We use zero-shot prediction of OTU dropout/colonisation between time points to demonstrate learned temporal stability at the week scale. Multiple samples for the same patient-timepoint are aggregated by union.

#### Snowmelt (zero-shot temporal seasonal) [23]

Alpine/cryosphere microbiome transitions across spring snowmelt, used to demonstrate learned temporal stability at the month scale. This dataset is particularly challenging as the communities being sampled are effectively exposed to the elements for long time periods between sampling. The various conditions included: compacted snow, ice encasement, ambient, no snow and no ice.

#### DIABIMMUNE (zero-shot temporal infancy-change) [24, 25]

Three-country infant cohort with repeated gut microbiome sampling at various time points during growth/infancy. This dataset is used to demonstrate learned temporal stability at long and short time scales, as well as the community-space attractor inference.

#### Infants (5CV) [26]

Infant gut cohorts with age (from birth to 5 years) and birth mode. Used for 5-fold joint age and birth mode classification to compare to MGM. Dataset preparation matched that described by MGM.

#### IBS (5CV) [27]

Curated multi-country populations for case-control classification and comparison to MGM [14]. This dataset consists of IBS and control subjects from three countries. Classification heads trained on one country are evaluated on others. Dataset preparation matched that described by MGM.

### 2.2 Representations

We embed bacterial 16S rRNA sequences using a DNA language model (ProkBERT) to obtain fixed-dimensional vectors without learned OTU IDs [20]. The full embedding space is shown in Fig. 2 We use ∼ 1 × 10^5^ SSU rRNA representative sequences (OTU level: 97%) from the MicrobeAtlas MAPseq reference database.

**Figure 2:**
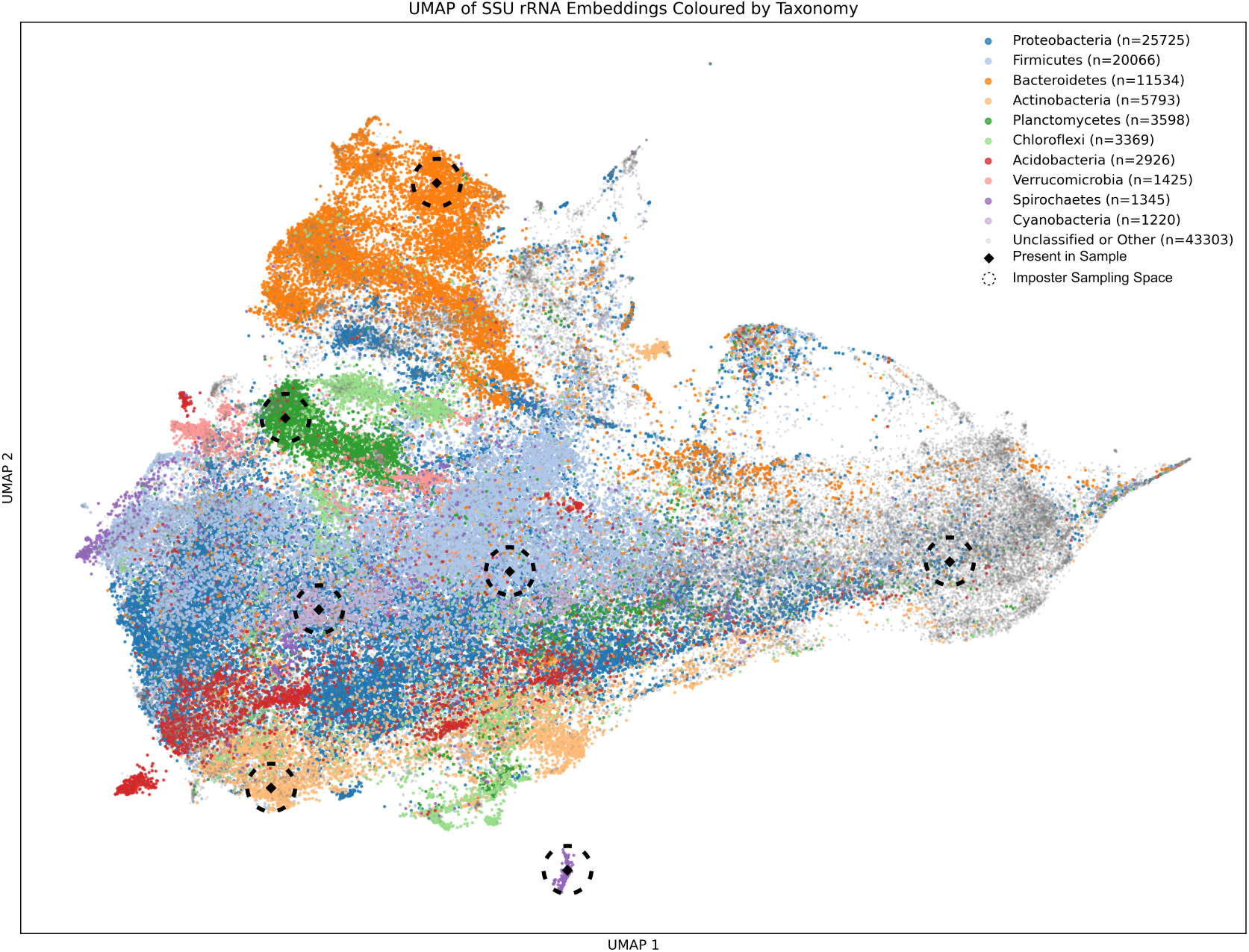
Imposter pool creation. Schematic of imposter selection in bacterial SSU-rRNA-embedding space and negative pool construction based on a sampled minimum cosine similarity value. Imposter OTUs are sampled from this imposter sampling space. This sampling strategy results in imposters of varying difficulty, sometimes even choosing plausible microbes as imposters.

We explored two variants of the denoising-set model:

(a) a **DNA-only** variant that only consumes SSU rRNA embeddings; and
(b) a **DNA + text** variant in which optional free-text metadata (biome, host, technology) are encoded with a large language model (text-embedding-3-small) and added as in-context tokens [28].

Unless explicitly noted, all main-text results are reported for the DNA-only variant, which performed slightly better on average across our evaluations.

Embedding space visualisations of DNA embeddings made use of UMAP reductions unless explicitly stated otherwise [29].

### 2.3 Negative generation

For each sample *x_i_* with observed OTUs *S_i_*, we construct a corrupted version 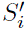 by adding a number of *imposter* OTUs that are relatively close to the sample in SSU-rRNA-embedding space (Fig. 2).

In practice, we first find a pool of minimum similarity OTUs for *S_i_* using cosine similarity on *ℓ*_2_-normalized SSU rRNA embeddings, and remove the OTUs already in *S_i_*. From this pool, we then sample roughly *r* · |*S_i_*| imposters (where *r* is a *noise rate*, i.e., the expected number of imposters per true taxon—in our case, 33%) and add them to the original set:

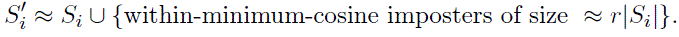

The minimum cosine cut off is also varied uniformly. This produces negatives that are often very similar to the true OTU, while still allowing variation in difficulty across samples. Metadata tokens, if provided, are left unchanged. More details on imposter sampling can be found in Supplementary Section S1.1.2.

### 2.4 Model and objective

For each sample *x_i_*, we start from a set of DNA embeddings *S^′^* = *S_i_* ∪ *N_i_* (true-present OTUs *S_i_* plus imposters *N_i_*) and the associated text embeddings *T_i_* (metadata, free text, etc. for text-trained model only).

#### Model

All embeddings are first mapped into a shared *d*-dimensional space with a simple type-specific linear layer.

We then feed the combined set into a Transformer encoder *T_θ_ without* positional encodings, so the model is permutation-invariant. This circumvents the requirement for arbitrary ordering of OTUs required by traditional Transformer based architectures. When present, text embeddings *T_i_* act as *sample-level context*: through self-attention they can influence the representations of all DNA embeddings. In the DNA-only variant we simply omit these tokens and train on the SSU rRNA embeddings alone.

A small output head *h_θ_*turns each DNA embedding into a scalar logit (stability score) that should be large for embeddings that were originally present, and small for imposters. We only interpret logits derived from DNA embeddings (*S^′^*), text tokens—when present—are used purely as context.

Full architecture specification and comparison between Susagi-Large and Susagi-Small are detailed in Supplementary Table S1.

#### Loss (DNA only, class-balanced)

We use a standard binary cross-entropy objective on logits and normalize positives and negatives separately so that both contribute equally.

#### Stability scoring

At inference time, a sample is again represented by *S* and *T*. The raw output of the model (logits) per SSU rRNA embedding are interpreted as member-level stability scores. A community stability score can be derived by simply averaging the logits of a sample *S*. We refer to the latter as the *community-level stability* score. In our setup, a higher score can be viewed as meaning more stable (in distribution, closer to training samples) and a lower score can be viewed as meaning less stable (out of distribution, further from training samples).

### 2.5 Model training

We use PyTorch ([30]) to train two denoising-set models (approximately 5×10^4^ and 8×10^5^ parameters) for two epochs on ∼ 2 × 10^6^ MicrobeAtlas samples using AdamW (learning rate 10*^−^*^4^), batch size 32, on a single V100 GPU. Training is performed once on MicrobeAtlas and the resulting encoder is reused for all downstream evaluations; we do not perform any additional fine-tuning.

### 2.6 Baselines and evaluation protocols

#### Zero-shot temporal prediction

For each longitudinal dataset we define per-OTU outcomes: *dropout* (present at *t*, absent at *t*+Δ) and *colonisation* (absent at *t*, present at *t*+Δ). Given the state *S_t_* (OTUs observed at time *t*, optionally augmented with text tokens), our pre-trained model produces per-OTU logits in a single forward pass, importantly with *no task-specific training*. For *dropout* this means simply passing the original sample through without alteration and reading off the logit on the OTUs of interest. For *colonisation* the readout is less trivial; as the OTU being tested is not present at *t*_1_, *S_t_*_1_ must be augmented by the inclusion of the OTU being tested so that it can have a logit from *t*_1_ that will be then be used to predict its appearance in *t*_2_.

We form all ordered *t*_1_→*t*_2_ pairs (chronology agnostic) within each grouping unit and evaluate by ranking OTUs using their logits, computing AUROC over OTUs among the samples. For induced gingivitis the grouping unit is *subject*; for DIABIMMUNE it is *subject*; and for Snowmelt it is *block–treatment* (e.g. C block, ice-covered).

#### Supervised temporal baselines

For comparison, we trained classical supervised baselines on the same tasks. Each sample at time *t* was represented as a binary presence/absence vector over OTUs, and the targets were per-OTU binary labels indicating dropout or colonisation between *t* and *t*+Δ. We trained one-vs-rest logistic regression, random forest classifiers, and a small multi-layer perceptron on these features using grouped 5-fold cross-validation to prevent leakage: grouping by *subject* for gingivitis and DIABIMMUNE, and by *block–treatment* for Snowmelt. For each task, we evaluated only OTUs for which both binary outcomes were observed in the data. For example, for dropout, an OTU was included only if there was at least one timepoint pair in which it was present at *t* and absent at *t*+Δ, and at least one pair in which it was present at both *t* and *t*+Δ. Analogous criteria were used for colonisation. We report mean ± s.d. macro-AUROC across folds.

#### 5-fold cross-validation (5CV) classification

For downstream classification tasks, we freeze the encoder and mean-pool token embeddings to form sample-level vectors, then train a linear classifier with stratified 5-fold CV. We evaluate on IBS vs control across countries, infant birth mode and age combined, and report AUROC.

We use MGM for comparison, following the author’s 5-fold cross-validation setup on classification tasks [14, 31]. Unlike MGM, we do not fine-tune the backbone models: we train only logistic regression heads on top of frozen sample embeddings. MGM serves as an autoregressive baseline operating on a fixed OTU vocabulary, whereas our model provides permutation-invariant SSU-rRNA-embedding-based features with fewer parameters overall.

#### Sample-level stability scoring within MicrobeAtlas

We compute stability scores for each sample and test enrichment of metadata keywords using Wilcoxon rank-sum with FDR, restricted to terms that appear at least 50 times (and with at least 50 samples without the term) after subsampling. The data was subsampled to ∼ 9 × 10^4^ samples uniformly for ease of computation. Susagi-Large (no-text) was used to generate the stability scores reported. Pielou’s evenness is calculated over raw abundance data to show that the model can infer abundance distribution from presence alone [32].

#### Community-space attractor visualisation

To probe attractor-like dynamics in community space, we performed a greedy anchored one-out-one-in rollout. At each step we updated the community by replacing the lowest-scoring microbe currently present with the highest of a uniformly sampled pool, if it scored better than the incumbent. An unchangeable anchor set was defined as community members that existed pre-rollout with model assigned probability over 98%. Each rollout was 300 steps long or cut short if there was no improvement in the mean score of the anchor set for 10 steps.

We fit PCA on the embedding vectors of real samples and projected both the real samples and rollout endpoints into the resulting 2D subspace. The community-space vector field was drawn by overlaying arrows from initial samples to rollout endpoints. The arrows were spatially coalesced by local grid-based averaging and scaled for visibility.

## 3 Results

### 3.1 The Imposter Denoising objective learns community structure

To illustrate how Susagi takes DNA embeddings and transforms them into presence/absence preditions, we create layer by layer visualisations of Susagi’s internal process.

UMAPs of token representations (gingivitis dataset) as they pass through Susagi show progressive separation of added vs. resident OTUs through (Fig. 3). Initially, imposters are interspersed in the SSU rRNA embedding space with the original OTUs of the sample. With each passing layer, the embeddings of the imposters and originals begin clustering in different areas of the model’s latent space. Finally, by layer 5, two very distinct clusters emerge representing the model’s prediction on wether those OTUs are original or not. More examples can be found in Supplementary Figure S2.

**Figure 3:**
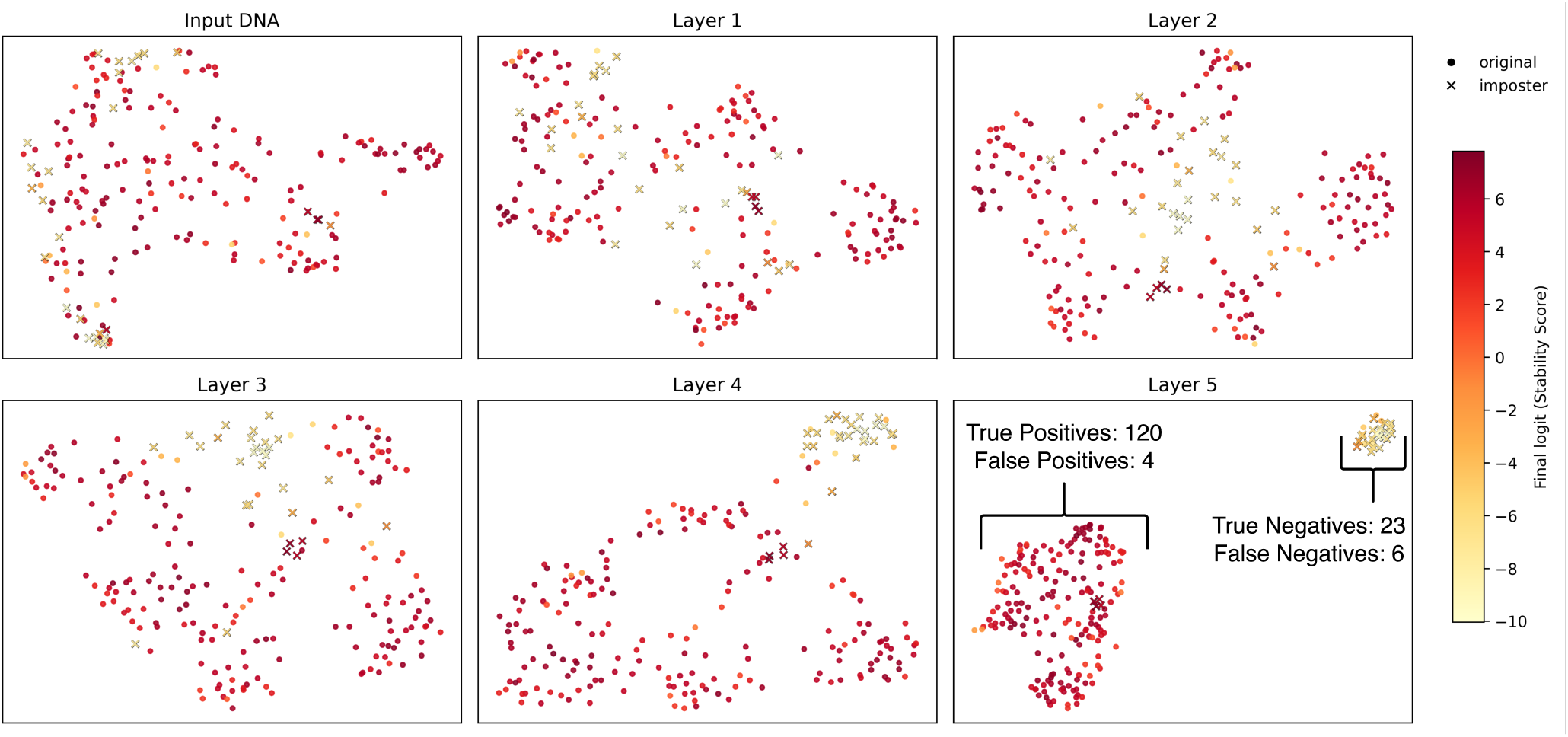
Layer-wise UMAPs illustrate imposter separation. Crosses: added OTUs (imposters); circles: resident OTUs; colour encodes final logits (per-OTU stability score). **Top left**: raw SSU-rRNA-embedding space. This representative sample was taken from the held-out Gingivitis dataset. For more examples see Supplementary Figure S2.

**Figure 4:**
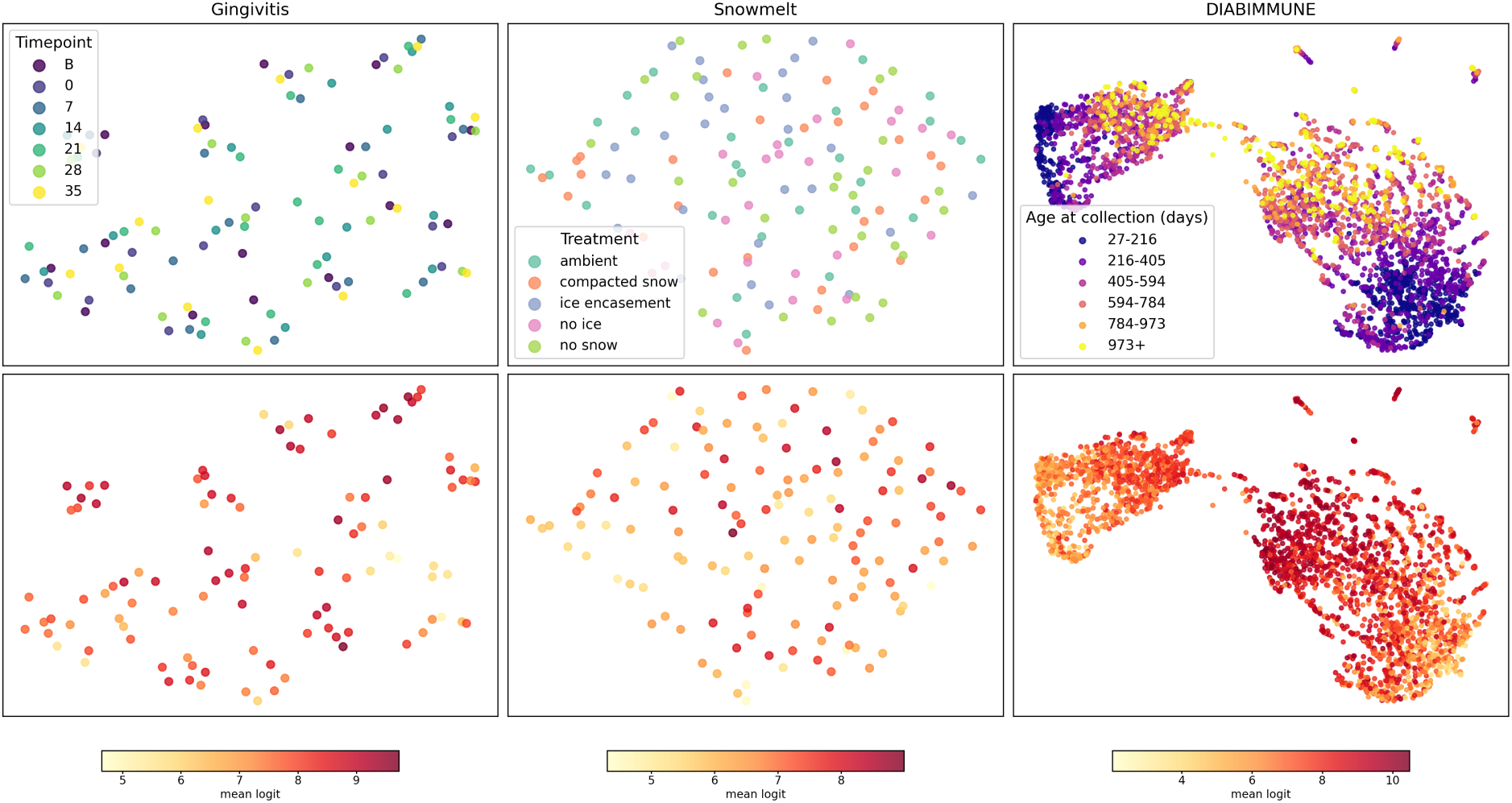
Dropout and colonisation benchmark dataset UMAPs. **Top row:** meta-data colouring on presence/absence Jaccard distances. **Bottom row:** sample stability score (mean logit) colouring from Susagi-Large (no-text). **DIABIMMUNE:** the sample stability score is strongly associated with subject age, and this age association is also evident in Fig. 6. **Gingivitis:** metadata colours indicate time point from baseline to day 35. **Snowmelt:**colours denote various treatments.

### 3.2 Time Series Evaluation

To understand whether Susagi’s logits encode a stability function over the SSU rRNA space, we devised a time-direction-agnostic dropout and colonisation task. If the model’s logits successfully correlate with the likelihood of an OTU being present at any arbitrary future or past time point, despite experimental perturbation, then they likely reflect a learned context-dependent stability function. Effectively, if an OTU receives a high logit, given a certain community, then it is expected not just to be present, but to be present for some time (resilient). If it is given a low logit, then it is expected to easily drop out again (transient).

To probe the model’s ability to generalise across varying contexts, we chose three datasets representing diverse environments and perturbations: oral (induced gingivitis), soil (alpine snowmelt), and infant gut. The varying time periods between sampling points and the various perturbations induced by the experimental protocols add another layer of difficulty to this task.

Without temporal supervision, Susagi-Large (no-text) predicts OTU dropout and colonisation above random and outperforms traditional ML models trained specifically for this task by 0.21 points on average (Fig. 5, Table. 1). Susagi-Large (no text) also surpasses the small variant on all datasets by 0.11 on average. Text training appears to be beneficial on certain datasets but only for the smaller models. Because we enforce strict grouped cross-validation by subject or block–treatment, and because OTU profiles can be sparse, per-study supervised baselines struggle to learn temporal structure that generalises across subjects, whereas Susagi (large and small) models can exploit the learned stability function from millions of samples. When cross-validation is relaxed, the performance of traditional ML improves notably (Supplementary Table S2).

**Figure 5:**
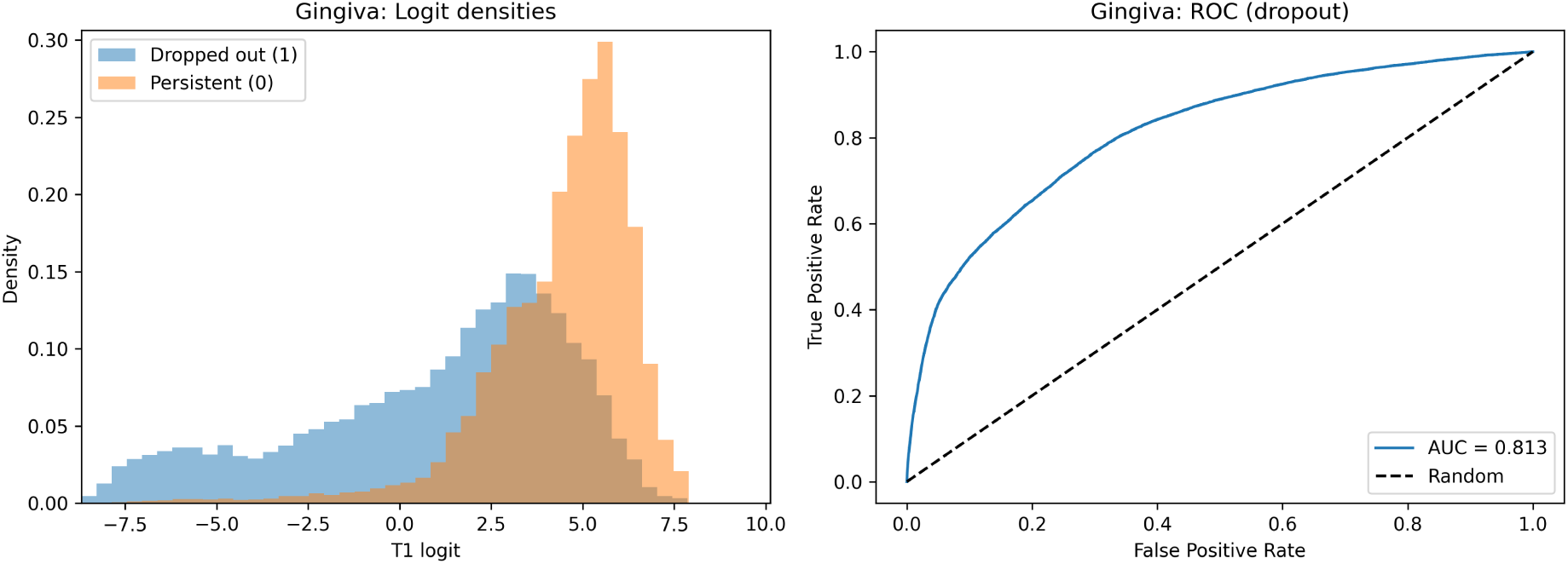
Zero-shot temporal dropout prediction in induced gingivitis. **Left:** dropout-prediction logit density plots, coloured by persistence (OTUs present at both time points) or dropout (OTUs present only at the first time point), showing that persistent OTUs are shifted toward higher logits while dropped-out OTUs are shifted toward lower logits. **Right:** ROC curve showing far-above-random performance. See Supplementary Fig. S3 for additional results.

**Table 1:** AUROC for OTU dropout (Dropout) and colonisation (Colonisation) across datasets (higher is better). Traditional ML methods shown: random forest, logistic regression, multilayer perceptron. Bold writing highlights the highest score. Numbers in dataset headers are the numbers of samples per dataset after sample grouping. Random baseline on any task is 0.5. ‘text’ and ‘no-text’ refer to models trained with metadata and without metadata. Traditional method results are 5-fold average AUCs.

| Model | Gingivitis (n=105) |  | Snowmelt (n=150) |  | DIABIMMUNE (n=3,348) |  |
| --- | --- | --- | --- | --- | --- | --- |
|  | *weeks–months* |  | *months–quarters* |  | *months–years* |  |
|  | Dropout | Colonisation | Dropout | Colonisation | Dropout | Colonisation |
| Susagi-Large ($\sim 8 \times 10^5$, no-text) | **0.81** | **0.71** | **0.65** | **0.58** | **0.68** | **0.82** |
| Susagi-Large ($\sim 8 \times 10^5$, text) | 0.79 | 0.70 | 0.58 | 0.55 | 0.66 | 0.76 |
| Susagi-Small ($\sim 5 \times 10^4$, no-text) | 0.65 | 0.66 | 0.50 | 0.51 | 0.60 | 0.70 |
| Susagi-Small ($\sim 5 \times 10^4$, text) | 0.67 | 0.64 | 0.50 | 0.56 | 0.56 | 0.62 |
| Random forest (500 trees) | $0.50 \pm 0.02$ | $0.50 \pm 0.01$ | $0.49 \pm 0.01$ | $0.51 \pm 0.01$ | $0.49 \pm 0.01$ | $0.53 \pm 0.03$ |
| Logistic regression (L2) | $0.50 \pm 0.02$ | $0.49 \pm 0.01$ | $0.49 \pm 0.02$ | $0.51 \pm 0.01$ | $0.48 \pm 0.01$ | $0.50 \pm 0.01$ |
| MLP (1 hidden, 128 dims) | $0.50 \pm 0.02$ | $0.48 \pm 0.02$ | $0.51 \pm 0.02$ | $0.50 \pm 0.01$ | $0.48 \pm 0.02$ | $0.50 \pm 0.02$ |

### 3.3 Classification Evaluation

#### Infants (unsupervised & 5CV)

In order to test whether the representations created by our model’s can have further downstream utility—outside of time series predictions—we tested the model on an infant birth mode and age prediction task. The Infants dataset consists of ∼ 2 × 10^3^ samples. Each sample is annotated with an age and birth mode, indicating the age at which the sample was taken, and the birth mode of the child (vaginal or caesarean).

UMAP of sample embeddings reveals age gradients (Fig. 6), recapitulating the pattern seen in Fig.4 and following well established developmental trends. The natural symmetry between the learned sample-level stability function and subject age likely bolstered the predictive performance of the frozen features in 5 cross-validation classification (Table 2). Our models use a simple linear regression head on top of frozen embeddings, whilst MGM employs fine-tuning of the underlying model weights. Again, Susagi-Large performed better than Susagi-Small (+0.09 accuracy, +0.03 mAUROC), indicating that the increased capacity of the model may have improved the quality of it’s representations. The performance of our large mode is very similar to MGM (∼ 0.55 accuracy, 0.91 mAUROC) suggesting this may be approaching the upper limit for the dataset.

**Figure 6:**
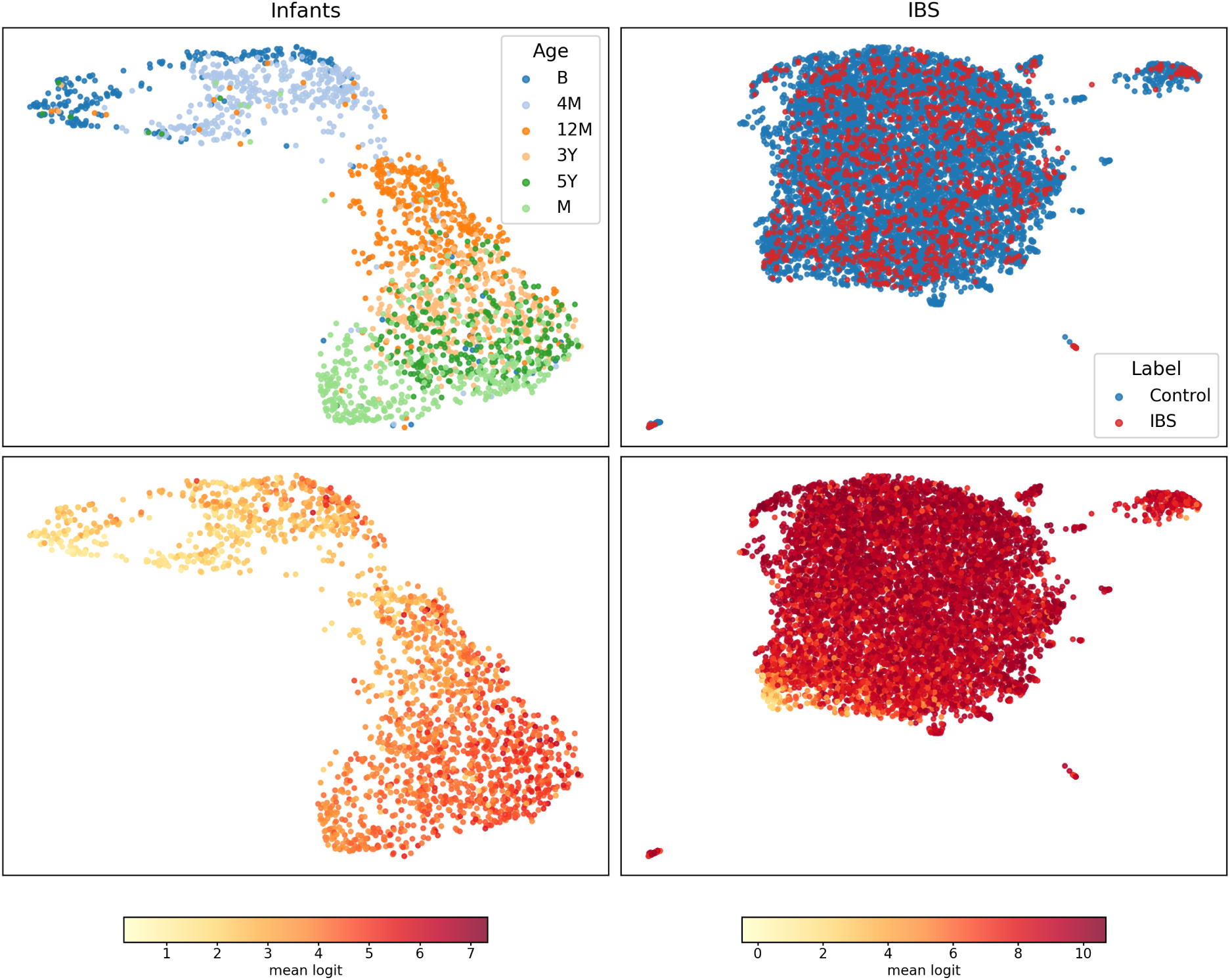
Classification task benchmark dataset UMAPs. **Top row:** metadata colouring on presence/absence Jaccard distances. **Bottom row:** sample stability score (mean logit) colouring from Susagi-Large (no-text). **Left to right:** datasets used for the classification tasks. **Infants:** metadata indicates age (birth to 5 years) and mother samples. As in Fig. 4, the stability score in the Infants dataset is strongly associated with age. **IBS:** metadata indicates control versus professionally diagnosed IBS.

**Table 2:**
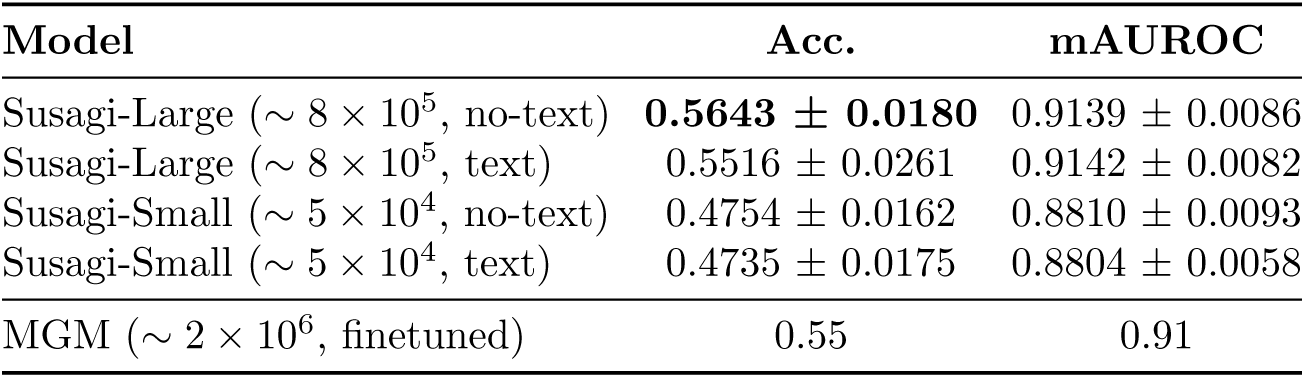
Accuracy and macro AUROC for joint age–delivery mode prediction on the Infants dataset using frozen unsupervised features with 5-fold cross-validation (higher is better). MGM had been fine-tuned on the underlying samples, whereas our models used frozen embeddings

#### IBS (5CV)

In this dataset the task involves predicting whether a subject has been diagnosed with IBS or is healthy.

Susagi’s frozen-encoder features are most competitive in the data-sparse regime (Table 3). When trained on the smallest source cohort (AUS, *n* = 303), our models outperform MGM across all test countries, with absolute AUROC gains of +0.06 to +0.26 (e.g., AUS→AUS: +0.19 for Large and +0.26 for Small). By contrast, MGM tends to have the edge when trained on the largest source cohort (USA), consistent with fine-tuning benefiting most when ample training data are available.

**Table 3:**
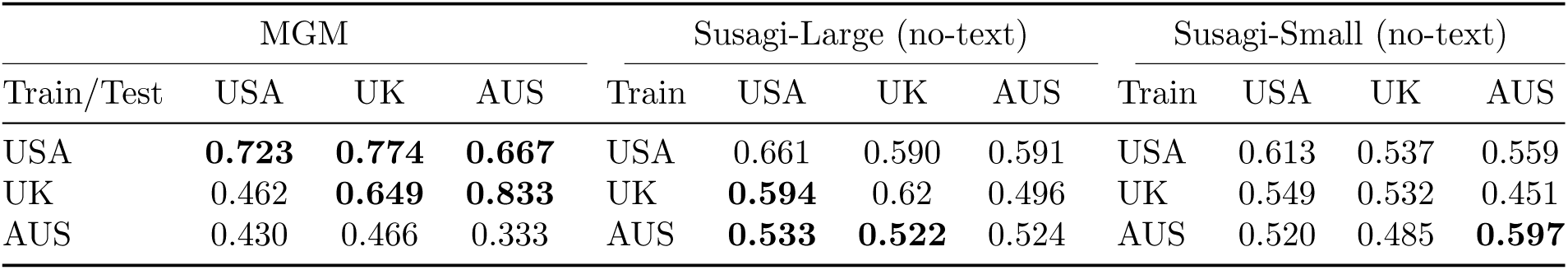
Cross-country AUROC for the fine-tuned MGM model and our fixed frozen embeddings (large and small). The highest AUROCs per train/test pair are in bold. The number of samples per country were: 303, 2294 and 5318 for AUS, UK and USA respectively.

### 3.4 Sample-level stability scoring in MicrobeAtlas

Although including metadata did not strongly affect model performance, predicted sample stability may still correlate with sample-associated annotations. To test this, we sampled 9 × 10^4^ microbiome samples from MicrobeAtlas spanning a wide range of environments and hosts, and derived a stability score for each sample.

Stability was strongly structured by annotation terms and biomass (Table 4): agricultural-, livestock-, and rumen-associated terms showed the largest positive effects on stability, whereas insect-associated terms and low-biomass communities (low OTU richness) showed the largest negative effects. However, the extended list (Supplementary Figure 7) indicates that the rela-tionship between biomass and stability is not always straightforward and generally has a weak correlation.

**Table 4:** Top eight enriched and depleted annotation terms ranked by effect size. Effect is the mean stability difference (with term − without term, logits). Susagi-Large (no-text) was used to generate the stability scores. Percentage indicates fraction of the sampled set with the term. Δ mean size represents the difference in the average number of OTUs (richness) for communities containing this term compared to background. For an extended list see Supplementary Figure S3.

| Top 8 enriched terms (higher stability) |  |  |  | Top 8 depleted terms (lower stability) |  |  |  |
| --- | --- | --- | --- | --- | --- | --- | --- |
| Term | Effect | $\Delta$ mean size | % with | Term | Effect | $\Delta$ mean size | % with |
| farmland | +1.55 | +587 | 0.1 | fungus | -1.46 | -242 | 0.1 |
| rumen | +1.49 | +267 | 0.9 | pitcher | -1.38 | -257 | 0.1 |
| earthworm | +1.46 | +578 | 0.1 | gill | -1.36 | -192 | 0.1 |
| cropland | +1.32 | +854 | 0.1 | lemur | -1.31 | -210 | 0.1 |
| arable | +1.27 | +481 | 0.1 | cell | -1.23 | -228 | 0.1 |
| scrofa | +1.23 | +157 | 0.2 | bumblebee | -1.18 | -279 | 0.1 |
| sus | +1.19 | +150 | 0.3 | virome | -1.17 | -236 | 0.1 |
| cow | +1.12 | +329 | 0.4 | fungi | -1.11 | -272 | 0.2 |

When analysing the relationship between stability score and OTU richness, the model generally followed a bell-shaped pattern in which the highest and lowest stability scores were assigned to the smallest communities (Supplementary Fig. 11). As community size increased, samples tended to receive stability scores closer to the average, typically slightly below 0, indicating that the average OTU had a probability of less than 50% of being present. For enriched terms, however, this relationship differed somewhat: communities associated with enriched terms could have larger community sizes and higher average stability scores than communities from stability-depleted environments. In fact, in order to achieve the highest stability scores, a sample would have to be both large and have high Pielou evenness (Figure 7). A random forest fit to predict stability score from evenness and richness explained 28% of the variance in the score in a uniform test split.

**Figure 7:**
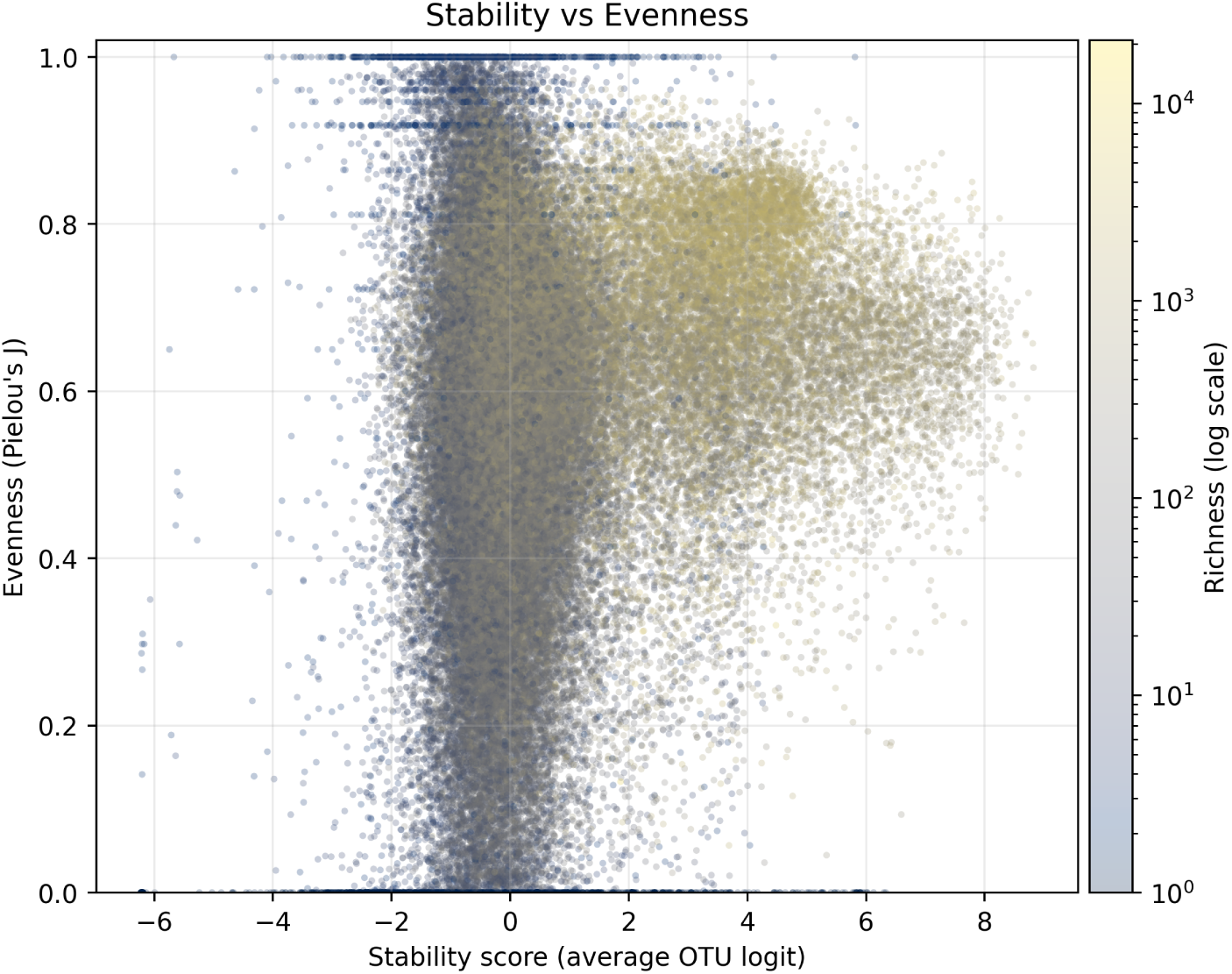
Stability versus community evenness, colored by community size (count). Evenness is computed as Pielou’s evenness using OTU abundances (the models never had access to this information). Points are samples, and colour indicates community size (count). The general pattern is that the highest stability scores are reached when samples are both highly even and large; having only one of these properties is typically not sufficient to produce the highest stability values.

### 3.5 Community-space Attractors

We hypothesise that one way the model makes temporal stability predictions is by relying on learned community-level priors, i.e., coarse “typical” profiles that can be robust to systematic experimental noise. Importantly, such priors could reflect true ecological structure but may also absorb dataset– or assay-specific regularities (for example, systematic differences between amplicon and WGS measurements). To explore this, we take real time points from the DIA-BIMMUNE dataset and perform a one-out-one-in simulation. The simulation works by defining an anchor set of highly stable members. At every step, the lowest-scoring non-anchor member is replaced by the highest-scoring member from a uniformly sampled candidate pool.

In Fig. 8 see that communities naturally converge to a common membership space attractor just beyond the most stable ends of the real data manifold. We theorise that this attractor represents a stable state for this community neighbourhood. Hence, predictions of community member stability are made based on this aggregate internal stereotype.

**Figure 8:**
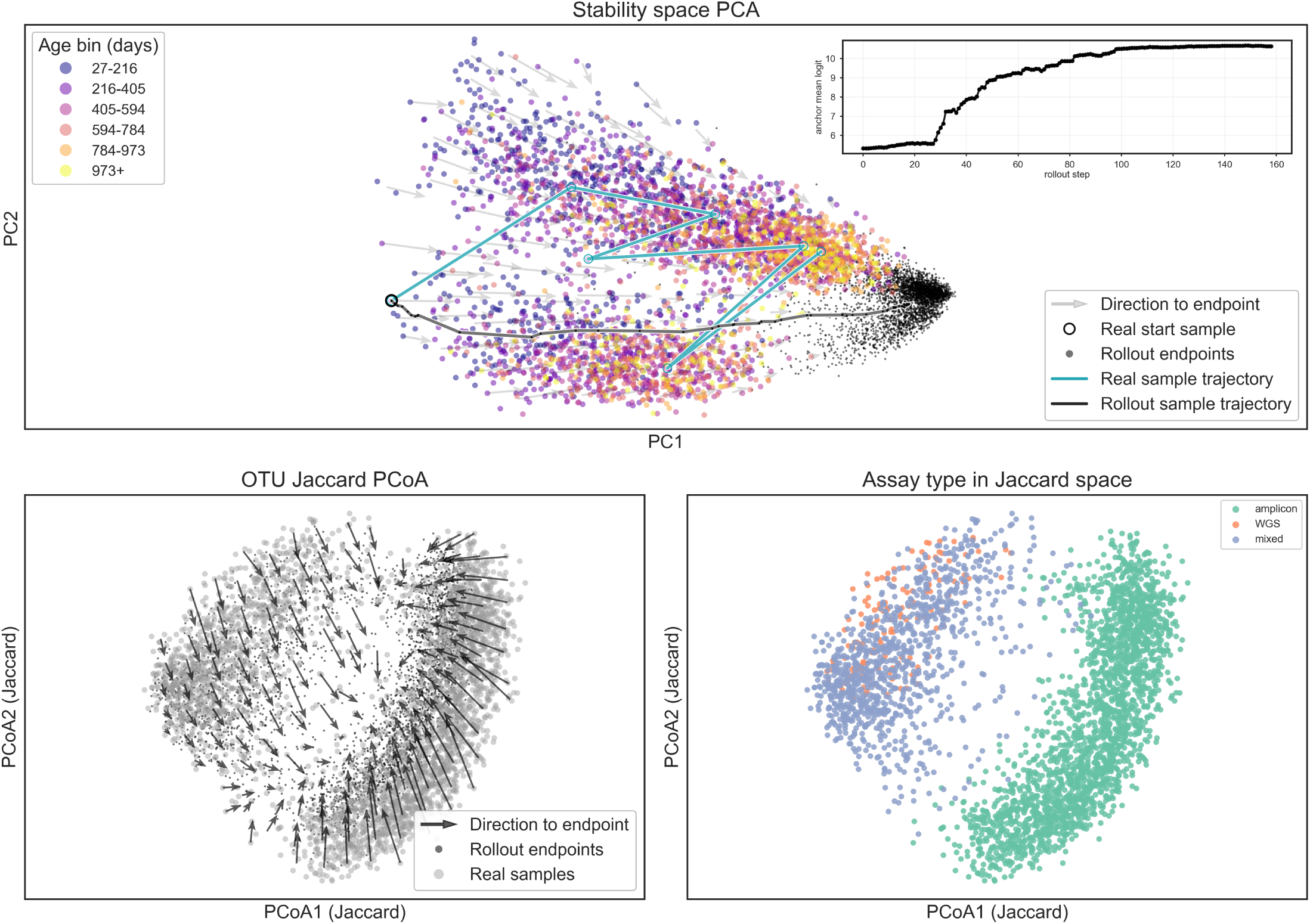
Overlay of one-out-one-in anchored rollout trajectories on the DIABIMMUNE embedding space. **Top**: PCA (PC1/PC2) of model embeddings for real samples (coloured by age bin) with rollout endpoints overlaid and arrows indicating displacement from each start point to its rollout endpoint; arrows are spatially coalesced by local grid-based averaging to reduce visual clutter. An inset shows the evolving stability of the sample’s anchor set through out the simulation. **Bottom-left**: 2D PCoA of OTU-set Jaccard distances with the same grid-averaged displacement arrows, real samples greyed for readability, and rollout endpoints shown in black. **Bottom-right**: Jaccard PCoA coordinates coloured by assay type (WGS/amplicon/mixed). “Mixed” indicates subject–age groups containing runs from more than one assay category.

Community trajectories (Fig. 8, top, black and turquoise lines) highlight an individual’s simulated and observed community progression with time. This shows that the increasing stereotypy of communities as individuals age is not a statistical artifact seen only at population level. This is similar to the trend seen in Fig. 4 and Fig. 6 where as individuals age, their microbial makeup tends toward the models learned stereotype in their community neighbourhood.

## 4 Discussion

We introduce a permutation-invariant denoising transformer that treats microbiome profiles as unordered sets of SSU rRNA sequence embeddings and learns a member-level notion of community stability. Trained on more than ∼ 2 × 10^6^ presence/absence 16S rRNA microbiome samples from MicrobeAtlas [5, 6], the model generalizes without training (zero-shot) to three longitudinal settings and provides reusable sample-level embeddings for downstream tasks. By operating directly in taxon-embedding space derived from MAPseq OTUs [33] and ProkBERT [20], the approach remains open-vocabulary and decouples community modelling from the choice of taxonomy or reference set.

### What the denoising objective learns

Our denoising objective is closer to contrastive learning than to standard supervised classification: instead of predicting labels or abundances per sample, the model learns to discriminate truly resident OTUs from nearby imposters in SSU-rRNA-embedding space, conditioned on the rest of the community. We interpret microbiome samples as approximate “stable states” of microbial communities, analogous to how large sequence models treat natural protein or genome sequences as outcomes of evolutionary selection [e.g. 34]. In protein and genomic language models, logits over single-residue or base substitutions correlate with fitness or structure despite training only on sequences; by analogy, training our model to recognise microbial communities as stable and reject local perturbations encourages it to internalize constraints that define community-level stability. Consistent with the stability interpretation, one-out-one-in anchored rollouts typically moved communities separated by sampling technology toward one another. The stable manifold appears to run between the two clusters and be slightly biased towards the higher age samples.

Thus, model-predicted logits can be viewed as context-dependent membership scores for OTU combinations, indicating which configurations are compatible with real, observed community states in MicrobeAtlas. Zero-shot temporal evaluations across induced gingivitis, alpine snowmelt, and DIABIMMUNE [22–25], spanning weeks to years, also support this interpretation: OTUs with high inferred residency in any given instant are more likely to persist across time, while low-scoring OTUs tend to drop out. Notably, the model infers these dynamics without temporal supervision, suggesting that zero-shot learning implicitly extracts predictive microbiome assembly roles with potential biological relevance. Traditional ML baselines trained directly on each temporal dataset underperform our zero-shot model, suggesting that the learned stability function is both non-trivial and transferable across settings, in line with broader experience that leveraging large unlabelled corpora improves microbiome prediction [12–14, 31]. At the same time, microbiome measurements are noisy and compositional [35], shaped by sampling depth, technical artefacts, detection limits, and contingent ecological events. Even an ideal model that captures the dominant stability constraints should therefore not be expected to achieve near-perfect AUROC, and performance must be interpreted against this noisy experimental reality.

Furthermore, stability scores derived from averaged sample-level logits suggest that much of the observed term-level enrichment and depletion reflects differences in how stability relates to community complexity across annotated sample groups. Positively enriched terms were dominated by agricultural and cropland annotations, consistent with prior work showing that soil microbiomes are highly diverse and that greater soil microbial diversity can support more stable ecosystem functioning [36]. Notably, the highest-scoring samples were characterised by both high Pielou evenness and large community size. This suggests that the model associates the most stable communities with high evenness, biomass and niche complexity, all things in common with the high stability enriched terms. Because the model has no access abundance data, this means the evenness of the distribution is being inferred purely by which microbes are present. In contrast, negatively enriched terms included placenta-, blood-, and insect-associated annotations, which are often regarded as low-biomass microbiome contexts where biological signal is weaker and more variable [37, 38].

### Comparison to autoregressive and taxonomy-centric models

Our design contrasts with autoregressive models such as MGM [14, 31], which impose an explicit ordering over OTUs (often by abundance) and model communities taxon-by-taxon over a fixed vocabulary of OTU IDs. Such autoregressive approaches are natural for sequence-like tasks (e.g. generation) and have shown strong performance on classification benchmarks [14], but they can entangle ordering artefacts with ecological structure and require retraining whenever the underlying vocabulary or taxonomic resolution changes. In contrast, our set-based model is explicitly permutation-invariant and consumes continuous SSU rRNA sequence embeddings rather than discrete OTU identifiers, allowing it to score previously unseen OTUs and remain valid under reference updates without modification provided the same DNA model is used.

Conceptually, the model operates in a DNA-embedding space that already encodes phylogenetic and, to some extent, functional information [19, 20, 39], rather than in a rigid OTU index space. This connects to work on permutation-invariant neural architectures for set-structured data [16, 17] and attention-based models more broadly [15]. It is also consistent with current practice in microbiome foundation models, which increasingly treat samples as sequences of tokens and leverage large unlabelled datasets for pretraining [12–14]. In this picture, our model can be viewed as learning a “DNA compatibility map” over communities rather than a co-occurrence map of fixed OTUs.

On disease benchmarks, frozen features from our encoder with simple linear heads are competitive with MGM on infant tasks [26] despite using presence/absence rather than abundances and keeping the backbone model fixed. On IBS [27], performance is data-regime dependent: our fixed-encoder models are strongest in the low-data setting, whereas MGM tends to have the edge as training data increase. It could be the case that our presence/absence based representations are more robust under the lower data regime, but miss the opportunity to learn from abundance based information that cuts through noise with more data. More broadly, our results suggest that permutation-invariant set models operating in sequence-embedding space are a useful complement to autoregressive, taxonomy-centric microbiome language models [12, 14].

### Practical implications

In practice, the model supports several use cases beyond the specific benchmarks we report. First, per-OTU logits enable sample-level quality control: OTUs that are consistently scored as non-resident given the rest of a community and its metadata may indicate contaminants or mapping artefacts. Second, the same logits support *in silico* perturbation analysis, for example by asking which candidate OTU the model expects to colonise or drop out when added to a given community, potentially guiding experimental consortia design and complementing dynamical approaches based on graph neural networks or recurrent architectures [9–11]. Third, sample embeddings derived from mean-pooled encoder states provide compact representations that can be reused across tasks, datasets, and modelling pipelines, reducing the need to retrain large models per study. Because these embeddings inherit information from both SSU rRNA sequence space and global, cross-environmental community structure [6], they can complement existing feature engineering based on taxonomies, pathways, or hand-crafted ecological indices, in line with the broader trend towards foundation models and transfer learning in microbiome analysis [12, 13].

### Limitations

Several limitations qualify our findings. First, we operate on SSU-rRNA-derived MAPseq 97% OTUs with presence/absence only [33], ignoring abundance and strain-level diversity. This is likely suboptimal for diseases where quantitative shifts, subtle compositional gradients, or specific strain combinations are important, as suggested by the IBS results and by compositional analyses of microbiome data [35].

Second, the MicrobeAtlas itself is heterogeneous: sequencing protocols, primer sets, and other experimental variables vary across contributing studies, as is common also in other large-scale resources [7, 8]. Although common reference mapping and the use of sequence embeddings (study agnostic coordinate system) mitigate some study specific bias [6], the model may still partially learn technical signatures, especially if these correlate with particular biomes or hosts. Consistent with this, we find that näıvely conditioning on available free-text metadata does not improve performance and slightly degrades it on average, presumably because many ostensibly sample-level fields are in practice noisy, missing, or inherited from project-level descriptions (though OTUs may also act as predictive proxies of metadata variables [40]). Third, our negative member construction (imposters) is purely sequence-based and proximity-driven: we enrich for close neighbours in SSU-rRNA-embedding space, which increases training difficulty but also means that the negative pool likely contains true positives (biologically plausible OTUs) and underweights mistakes involving more distant OTUs. This can bias the learned notion of “stability” toward local neighbourhoods of DNA space defined by the underlying encoder [20]. Finally, our approach inherits the resolution limits of the upstream DNA and text embeddings [20, 28]. If the SSU rRNA encoder effectively collapses a region of sequence space into a single cluster, the model cannot distinguish OTUs within that region, regardless of data volume. Similarly, if coarse or noisy metadata are embedded into overlapping regions, this could blur important contextual differences. In both cases, the community-stability function we learn is bounded by what the underlying embedding spaces can resolve.

### Opportunities for extension

A central opportunity is richer conditioning, but the main bottleneck is the granularity and reliability of publicly available metadata: intervention and host-factor labels are often recorded at the project level rather than per sample, limiting the utility of text conditioning in large repositories [6, 8]. If high-quality structured sample-level metadata were available, the same architecture could support counterfactual queries (e.g. adding an antibiotic course to a baseline sample and inspecting per-OTU logit shifts), targeted community design (suggesting resilient or easily displaced members under specified conditions), and planetary-scale out-of-distribution detection across biomes and studies.

Overall, our results support the view that permutation-invariant denoising over SSU rRNA embeddings [16, 17, 20] is a viable way to learn reusable community-level representations and member-level stability scores, complementing taxonomy-centric microbiome language models [12–14].

## Data and Code Availability

Training corpus: unified dataset of SSU rRNA samples with metadata downloadable from https://microbeatlas.org/landing. Evaluation data: experimental gingivitis, snowmelt, infant cohorts, DIABIMMUNE, IBS can be found as part of the data package linked in the code repository. Code: public release under an opensource license https://github.com/the-puzzler/Microbiome-Modelling. Model weights can also be downloaded directly from https://huggingface.co/basilboy/microbiome-model. Interactive demo can be found here: https://huggingface.co/spaces/basilboy/microbiome-space.

## Funding and Acknowledgements

This work was supported by the University of Zurich and by the Swiss National Science Foundation project grant 310030 192569. The authors would also like to acknowledge the members of the von Mering group for their discussions on ideas and especially their feedback on the figures.

## S1 Supplementary Information

### S1.1 Model and training

#### S1.1.1 Model Architectures

The large and small versions of the model vary by embedding dimension, feedforward dimension size, number of layers and total number of parameters:

**Supplementary Table S1:** Model Hyperparameters.

| Model | $D_{\text{model}}$ | $N_{\text{head}}$ | Layers | Dropout | $\text{dim}_{\text{ff}}$ | OTU Emb | TXT Emb | Params (approx.) |
| --- | --- | --- | --- | --- | --- | --- | --- | --- |
| Small | 20 | 5 | 3 | 0.1 | 80 | 384 | 1536 | $\sim 50\text{k}$ |
| Large | 100 | 5 | 5 | 0.1 | 400 | 384 | 1536 | $\sim 800\text{k}$ |

The embedding dimensions of ProkBERT and text-embedding-3 are 384 and 1536 respectively. The total number of unique OTU embeddings was ∼ 1 × 10^5^.

#### S1.1.2 Imposter Sampling

In practice, rather than thresholding on a fixed cosine similarity, we first sample a value (k) uniformly between 2,000 and 90,000 and construct, for each sample, a candidate pool of the (k) most similar OTUs that are not already present. We then draw a subset of this pool as imposters (negatives), with the number of imposters scaled to the number of OTUs present in the original sample.

Without this method, it is very unlikely that very challenging examples would emerge by chance due to the huge space of possible embeddings relative to the number of samples. This constraint forces some examples to contain purely similar embeddings, thus training the model to be more cautious with decision boundaries in embedding space.

For training all model’s we used a 33% noise rate. This means that for a given sample being passed through the model, approximatly 33% of the OTU embeddings are not from the original sample.

#### S1.1.3 Inference Peculiarity

Whilst testing the model on downstream tasks, it was noted that the addition of “scratch tokens” appeared to generally improve the performance of the models “for free” (Fig. S1). Scratch tokens in this case are near zero vectors that are appended to the input sequence of 16S embeddings.

We theorise that this phenomena, that has been observed in other work, is likely linked to message passing between transformer layers [41]. During training, the model probably chooses low information content embeddings to store and pass information. However, with the addition of completely useless embeddings, this process is made more efficient, perhaps recovering some over-written information.

**Supplementary Figure S1:**
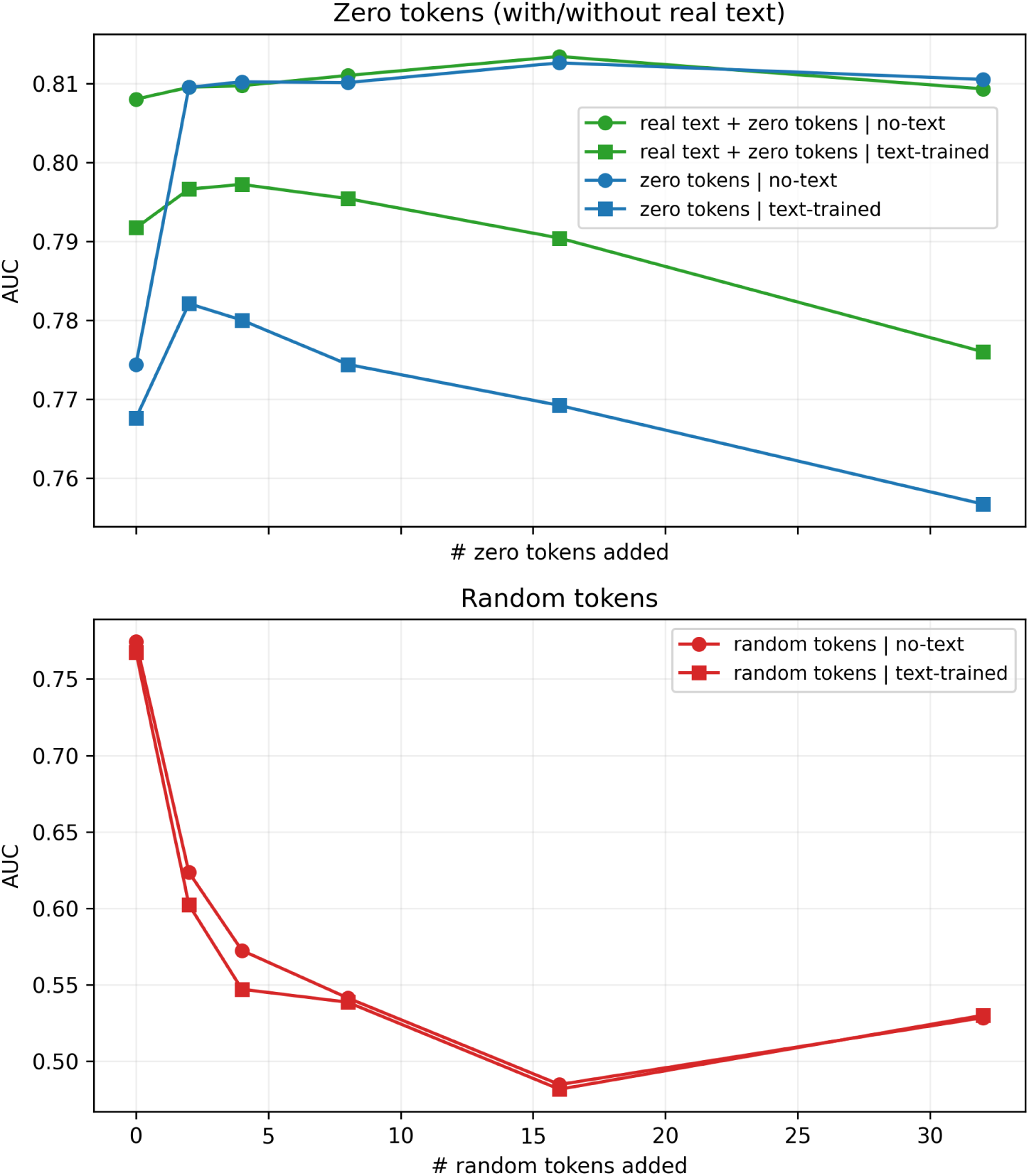
Gingivitis dropout AUC as a function of added tokens. Circles denote the no-text checkpoint, squares denote the text-trained checkpoint. **Top:** performance of no-text and text-trained models with the addition of zero tokens and/or meta-data derived (real text) tokens into model context. **Bottom:** no-text and text-trained models’ performance when randomly initialised tokens are added into context.

##### Observations

The no-text and text-trained models have different optimal numbers of scratch tokens. The no-text model is more robust: its performance improves and then plateaus as more scratch tokens are added, without clear degradation at higher counts. In contrast, the text-trained model peaks with a small number of scratch tokens and then declines as more are added. When real text is present, the no-text model performs about as well as when text is absent but scratch tokens are added, suggesting the added tokens are acting as a scratchpad rather than contributing semantic content.

Random tokens, however, quickly degrade both models (bottom panel). A likely reason is that the random tokens follow a very different distribution than the near-zero scratch tokens. For example, the text input projection in the no-text checkpoint has very small scale (mean ≈ −3.5×10*^−^*^5^, std ≈ 1.47×10*^−^*^2^), whereas the text-trained checkpoint has much larger spread (std ≈ 4.04 × 10*^−^*^2^, min/max ≈ −0.227*/*0.215), indicating that injecting random tokens introduces significantly higher-magnitude activations that disrupt inference.

### S1.2 Supplementary Results

#### S1.2.1 Internal layer representations of the no-text model

These visualisations (Fig. S2) are made by looking at the internal representations of the input DNA sequences and how they are transformed as they pass through the model.

The internal representations of the model allow us to gain some intuition behind its inner workings. The input DNA space map is manipulated with every layer, slowly transforming the space until a clear decision boundary appears.

**Supplementary Figure S2:**
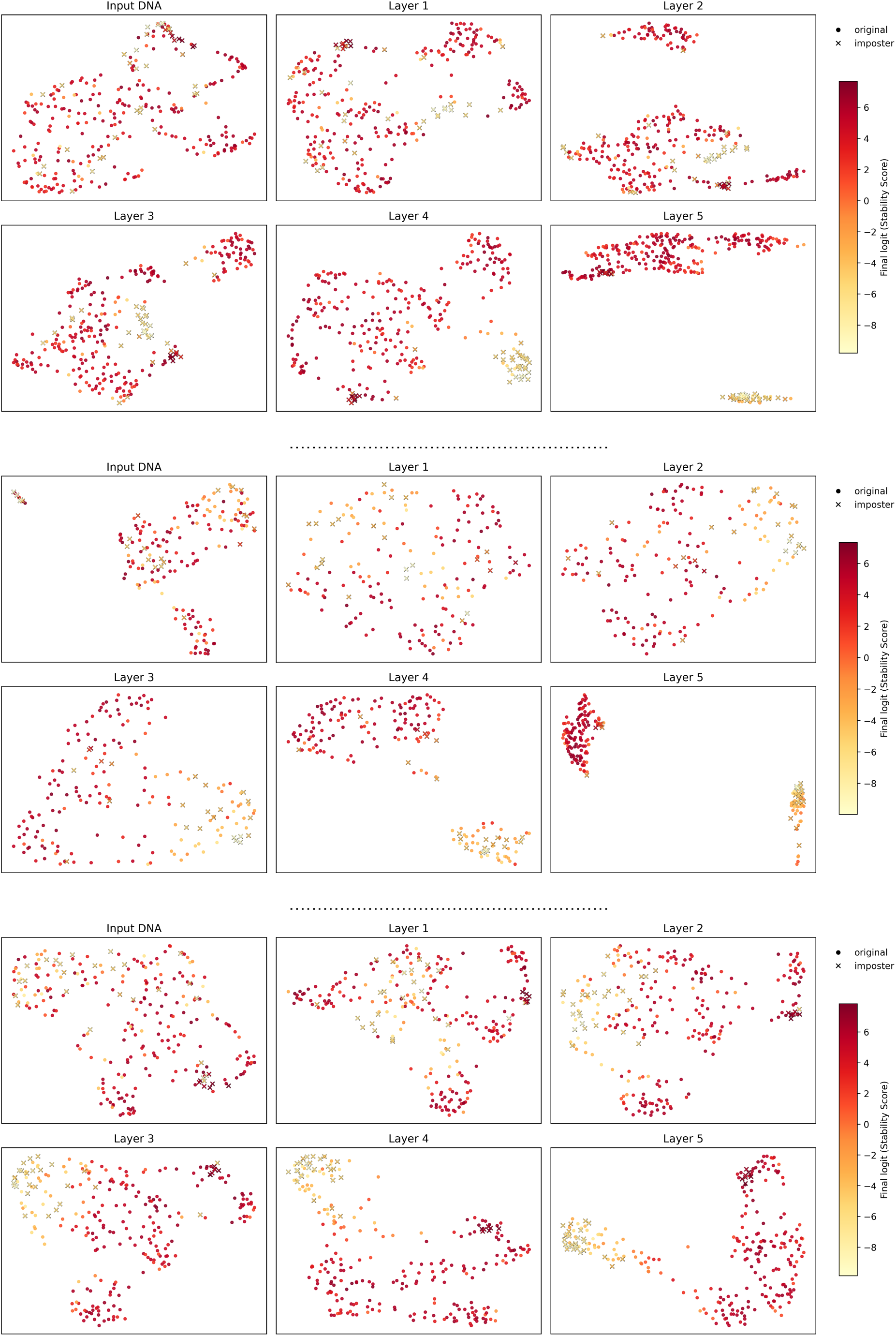
Layer-wise UMAPs for Susagi-Large (no-text) model. Crosses: added OTUs (imposters); circles: resident OTUs; colour encodes final logits (per OTU stability score). This figure shows more randomly chosen samples from the held-out Gingivitis dataset

#### S1.2.2 Additional time series ROC curves for the no-text model

**Supplementary Figure S3:**
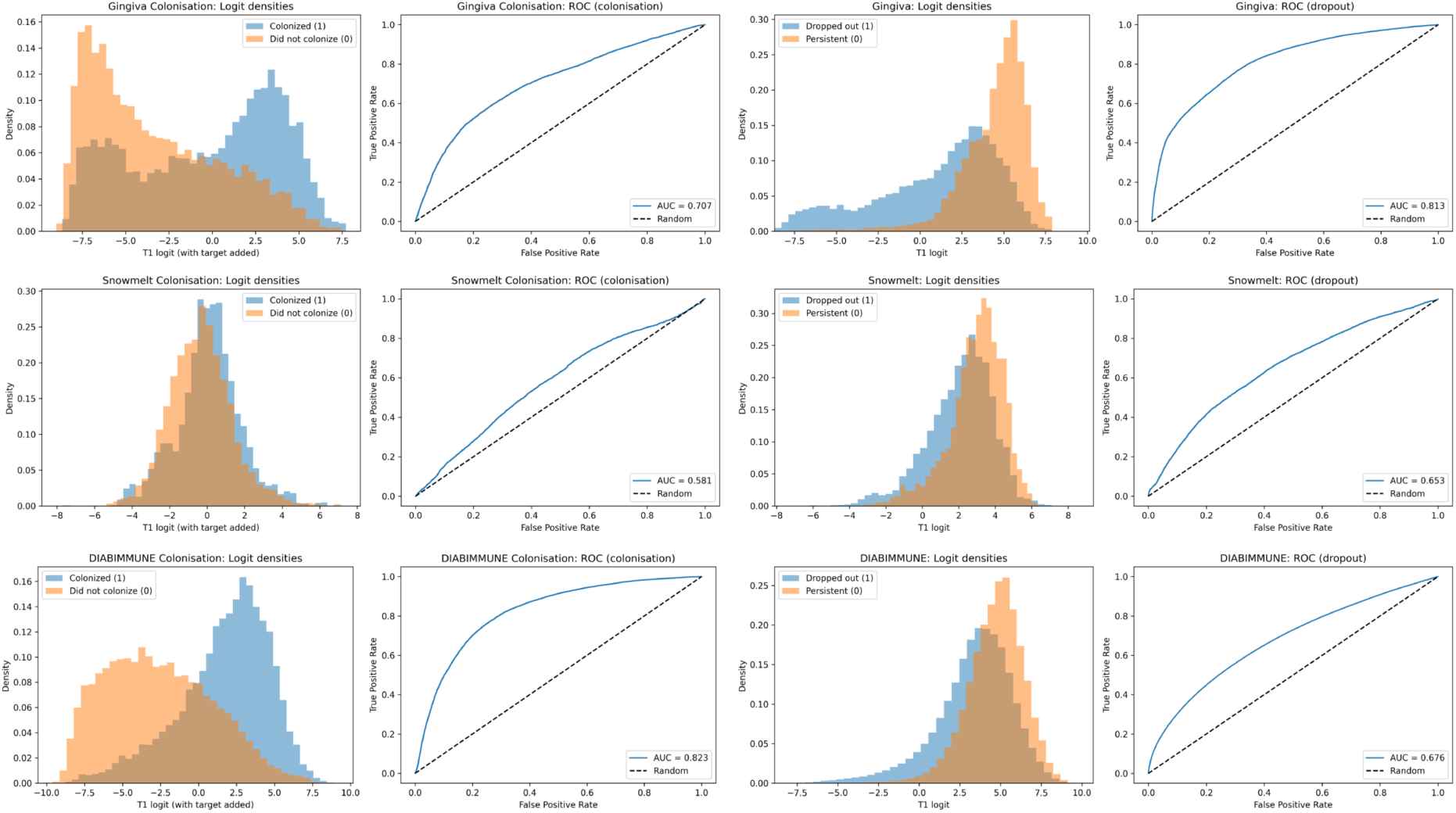
Zero-shot temporal dropout and colonisation prediction in Gingivitis, Snowmelt and DIABIMMUNE datasets. The left columns shows colonisation results and the right columns, dropout results.

#### S1.2.3 Relaxed Cross-Validation Baselines

**Supplementary Table S2:** AUROC for OTU dropout (Dropout) and colonisation (Colonisation) across datasets (higher is better) using a single random 80/20 split. Bold writing highlights the highest score per dataset/task. Random baseline on any task is 0.5. Numbers in dataset headers are the numbers of samples per dataset after sample grouping.

|  | Gingivitis (n=105) |  | Snowmelt (n=150) |  | DIABIMMUNE (n=3,348) |  |
| --- | --- | --- | --- | --- | --- | --- |
|  | *weeks-months* |  | *months-quarters* |  | *months-years* |  |
| Model | Dropout | Colonisation | Dropout | Colonisation | Dropout | Colonisation |
| Logistic regression (OvR) | 0.670 | **0.689** | 0.513 | **0.507** | 0.542 | **0.701** |
| MLP (1 hidden, 128 dims) | **0.675** | 0.685 | **0.541** | 0.494 | **0.553** | 0.699 |

#### S1.2.4 OTU Richness and stability score relationship

**Supplementary Figure S4:**
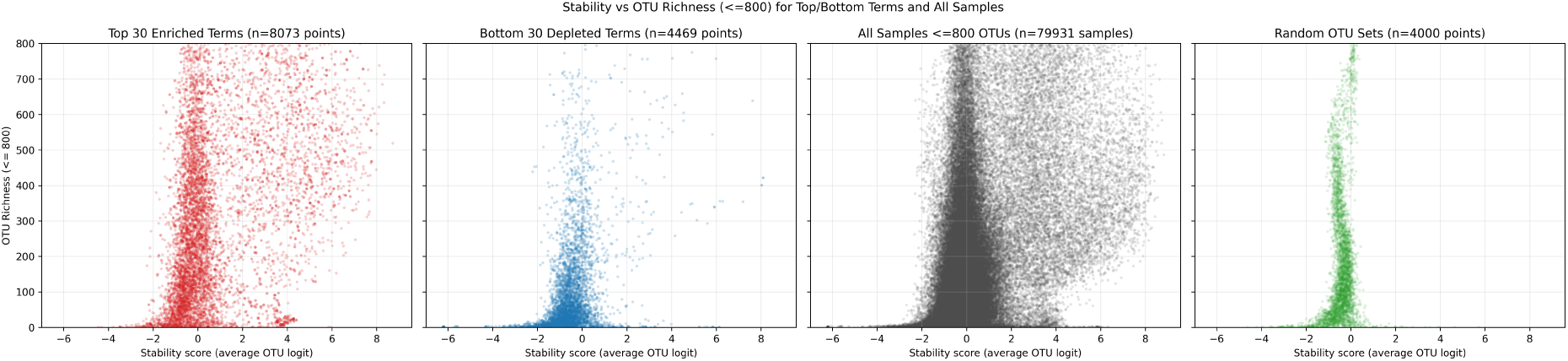
OTU richness and stability score relationships for the top 30 enriched and depleted terms. The enriched terms exhibit a right shifted relationship indicating that these terms support higher stability scores across a range of community sizes. Meanwhile, the depleted terms generally do not, and favour a left shifted bell curve shape more strictly. Communities composed of random OTUs are also left shifted with nearly no communities achieving over 50% average probability (0 score). For viewability, samples with more than 800 OTUs are not plotted.

#### S1.2.5 Extended keyword list and stability scores

**Supplementary Table S3:**
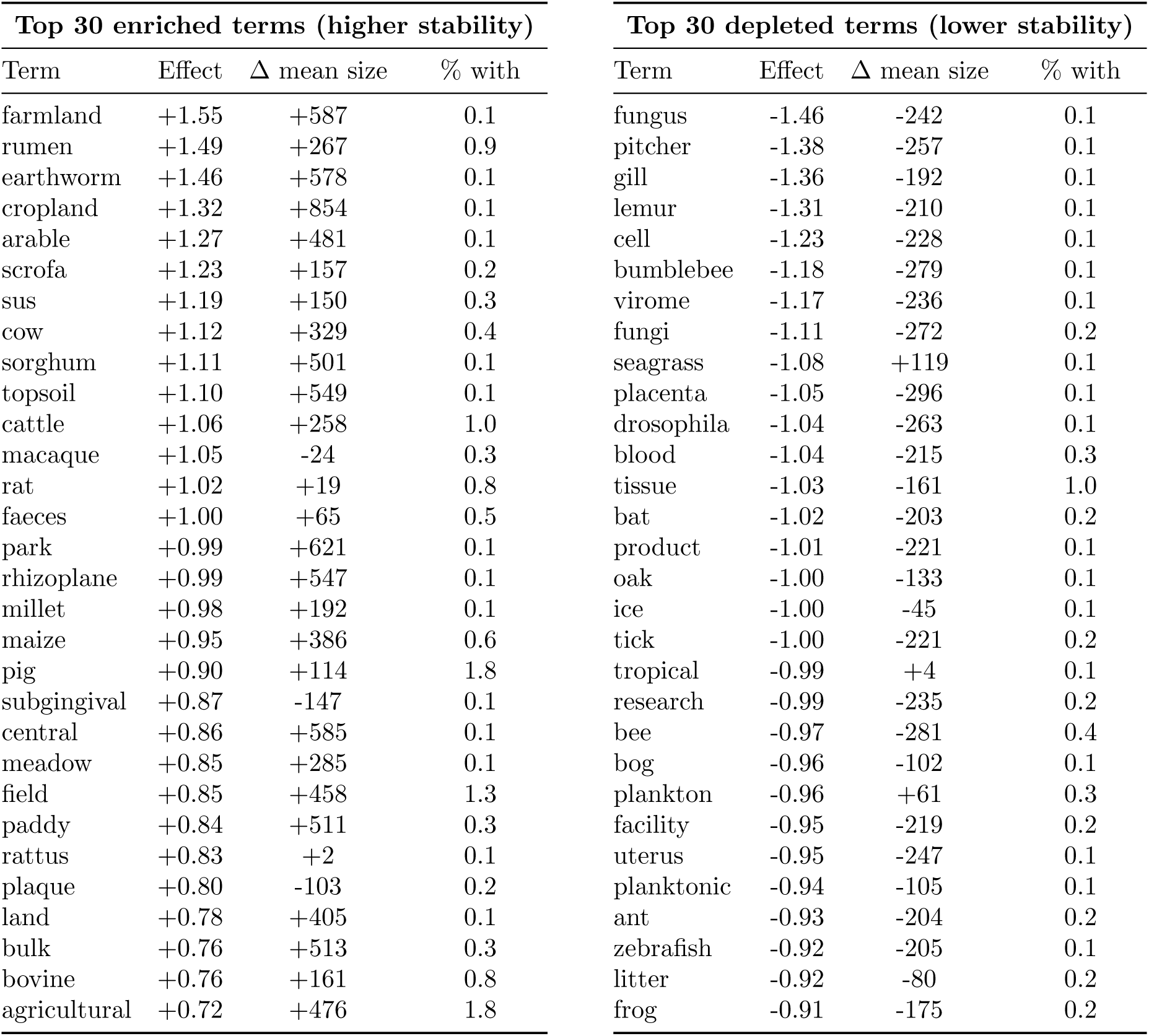
Top thirty enriched and depleted annotation terms ranked by effect size (mean logit difference, with term − without term, Susagi-Large no-text). Δ mean size represents the difference in the average number of OTUs for communities containing this term compared to background.

| Top 30 enriched terms (higher stability) |  |  |  | Top 30 depleted terms (lower stability) |  |  |  |
| --- | --- | --- | --- | --- | --- | --- | --- |
| Term | Effect | $\Delta$ mean size | % with | Term | Effect | $\Delta$ mean size | % with |
| farmland | +1.55 | +587 | 0.1 | fungus | -1.46 | -242 | 0.1 |
| rumen | +1.49 | +267 | 0.9 | pitcher | -1.38 | -257 | 0.1 |
| earthworm | +1.46 | +578 | 0.1 | gill | -1.36 | -192 | 0.1 |
| cropland | +1.32 | +854 | 0.1 | lemur | -1.31 | -210 | 0.1 |
| arable | +1.27 | +481 | 0.1 | cell | -1.23 | -228 | 0.1 |
| scrofa | +1.23 | +157 | 0.2 | bumblebee | -1.18 | -279 | 0.1 |
| sus | +1.19 | +150 | 0.3 | virome | -1.17 | -236 | 0.1 |
| cow | +1.12 | +329 | 0.4 | fungi | -1.11 | -272 | 0.2 |
| sorghum | +1.11 | +501 | 0.1 | seagrass | -1.08 | +119 | 0.1 |
| topsoil | +1.10 | +549 | 0.1 | placenta | -1.05 | -296 | 0.1 |
| cattle | +1.06 | +258 | 1.0 | drosophila | -1.04 | -263 | 0.1 |
| macaque | +1.05 | -24 | 0.3 | blood | -1.04 | -215 | 0.3 |
| rat | +1.02 | +19 | 0.8 | tissue | -1.03 | -161 | 1.0 |
| faeces | +1.00 | +65 | 0.5 | bat | -1.02 | -203 | 0.2 |
| park | +0.99 | +621 | 0.1 | product | -1.01 | -221 | 0.1 |
| rhizoplane | +0.99 | +547 | 0.1 | oak | -1.00 | -133 | 0.1 |
| millet | +0.98 | +192 | 0.1 | ice | -1.00 | -45 | 0.1 |
| maize | +0.95 | +386 | 0.6 | tick | -1.00 | -221 | 0.2 |
| pig | +0.90 | +114 | 1.8 | tropical | -0.99 | +4 | 0.1 |
| subgingival | +0.87 | -147 | 0.1 | research | -0.99 | -235 | 0.2 |
| central | +0.86 | +585 | 0.1 | bee | -0.97 | -281 | 0.4 |
| meadow | +0.85 | +285 | 0.1 | bog | -0.96 | -102 | 0.1 |
| field | +0.85 | +458 | 1.3 | plankton | -0.96 | +61 | 0.3 |
| paddy | +0.84 | +511 | 0.3 | facility | -0.95 | -219 | 0.2 |
| rattus | +0.83 | +2 | 0.1 | uterus | -0.95 | -247 | 0.1 |
| plaque | +0.80 | -103 | 0.2 | planktonic | -0.94 | -105 | 0.1 |
| land | +0.78 | +405 | 0.1 | ant | -0.93 | -204 | 0.2 |
| bulk | +0.76 | +513 | 0.3 | zebrafish | -0.92 | -205 | 0.1 |
| bovine | +0.76 | +161 | 0.8 | litter | -0.92 | -80 | 0.2 |
| agricultural | +0.72 | +476 | 1.8 | frog | -0.91 | -175 | 0.2 |

